# Transcriptional response of a target plant to benzoxazinoid and diterpene allelochemicals highlights commonalities in detoxification

**DOI:** 10.1101/2022.02.21.480921

**Authors:** Eva Knoch, Judit Kovács, Sebastian Deiber, Reshi Shanmuganathan, Núria Serra Serra, Claude Becker, Niklas Schandry

## Abstract

Plants growing in proximity to other plants are exposed to a variety of metabolites that these neighbors release into the environment. Some species produce allelochemicals to inhibit growth of neighboring plants, which in turn have evolved ways to detoxify these compounds. In order to understand how the allelochemical-receiving target plants respond to chemically diverse compounds, we performed whole-genome transcriptome analysis of *Arabidopsis thaliana* exposed to either the benzoxazinoid derivative 2-amino-3H-phenoxazin-3-one (APO) or momilactone B. These two allelochemicals belong to two very different compound classes, benzoxazinoids and diterpenes, respectively, produced by different cereal crop species. Despite their distinct chemical nature, we observed similar molecular responses of *A. thaliana* to these allelochemicals. In particular, many of the same or closely related genes belonging to the three-phase detoxification pathway were upregulated in both treatments. Further, we observed an overlap between genes upregulated by allelochemicals and those involved in herbicide detoxification. Our findings highlight the overlap in the transcriptional response of a target plant to natural and synthetic phytotoxic compounds and illustrate how herbicide resistance could arise via pathways involved in plant-plant interaction.

## Introduction

In the competition for nutrients and space, some plants secrete growth-inhibiting chemicals into the soil to inhibit the germination or growth of neighboring plants [1, 2]. These phytotoxic compounds, known as allelochemicals, can be released from the roots of the donor plant or leach into the soil from leaves or decaying plant material. Several species of the *Poales*, the economically most important order of plants, produce such allelochemicals, but their chemical nature can be quite diverse. For example, maize, wheat, and rye all produce benzoxazinoids (BX) [3–5], while many rice cultivars produce diterpenes such as momilactone B [6–8]. There is currently no example of a species that is capable of producing both momilactone B and BX compounds. This might suggest that - despite their very different chemical characteristics - these two types of allelochemicals exert redundant activities on target plants.

The two main forms of BX in cereals are DIBOA (2,4-dihydroxy-2H-1,4-benzoxazin-3(4H)-one) and its C-7-methoxy derivative DIMBOA, for which the biosynthetic pathways have been fully characterized (reviewed in [3]). BXs enter the soil either via exudation from the plant roots or via decomposing plant material that is incorporated into the soil. In an agricultural context, this can happen for example when grasses are used as cover crop or when straw is left on the field post-harvest as green manure [9]. In the soil, microbes rapidly degrade DIBOA to the final, stable product 2-amino-3H-phenoxazin-3-one (APO) [10]. APO and the analogous, DIMBOA-derived AMPO are potent phytotoxins [11] that inhibit histone deacetylases and slow down root growth [12].

As alluded to above, rice produces a very different class of allelochemicals than wheat, maize, and rye. Upon environmental cues, ranging from neighbor proximity to fungal attack to abiotic stress, rice plants produce the diterpenes momilactone A and B (reviewed in [13]). Momilactone A contributes to resistance to fungal pathogens [14, 15], while momilactone B has stronger allelochemical properties, inhibiting germination and root growth of a broad range of target species [16]. The biosynthesis of momilactone B has been fully elucidated [17–20]. Like DIBOA and DIMBOA, momilactone B is exuded from the roots by a yet unknown mechanism. Although its growth-inhibitory properties are well known, its mode of action remains unresolved [13].

As some plants release allelochemicals to inhibit growth of neighboring plants, their neighbors in turn have evolved ways to detoxify these compounds. As for xenobiotics such as herbicides, allelochemicals taken up by the target plant may be detoxified through a three-phase detoxification system [21, 22], by which the compounds are metabolically activated (phase I) to allow conjugation with sugars or amino acids (phase II). Soluble conjugates may then be transported to and stored in the vacuole or in the apoplast (phase III). Different classes of enzymes and proteins contribute to the different stages of detoxification: cytochrome P450s (CYPs), hydroxylases, and peroxidases activate xenobiotics in phase I, while phase II-detoxifying enzymes include UDP-dependent glucosyltransferases (UGTs), glutathione-S-transferases (GSTs), and quinone oxidoreductases [23]. Membrane transporters such as ATP-binding cassette (ABC), multi-antimicrobial extrusion (MATE) and major facilitator superfamily (MFS) transporters then remove the conjugates from the cytosol in phase III by sequestering them either in the vacuole or exporting them into the apoplast [24].

Here, we asked whether chemically distinct allelochemicals from different donor species but with similar growth arrest effects and potency trigger similar molecular responses in the target plant. We explored the transcriptional response of the model plant *A. thaliana* to two different and agriculturally relevant allelochemicals, the benzoxazinoid APO and the diterpene momilactone B. We profiled the immediate, short-term transcriptional response of *A. thaliana* seedlings to half-maximal effect concentrations of either compound by time series mRNA sequencing (RNA-seq). We hypothesized that by determining the commonalities and differences in molecular response to these two compound classes, we might be able to discern their respective mode of action and shed light on coping mechanisms of the target plant. The respective transcriptional response to the two allelochemicals showed substantial functional overlap. In particular, we identified components of the three-phase detoxification pathway to be up-regulated in both treatments. This suggests that, despite their distinct chemical characteristics, both types of allelochemicals trigger similar xenobiotic detoxification responses in the target plant, and that the involved enzymes have promiscuous functionality in detoxifying xenobiotics and other phytotoxic compounds.

## Results

### Different allelochemicals trigger similar transcriptional activation

To determine the transcriptional response of *A. thaliana* to the two allelochemicals APO and momilactone B, we exposed *A. thaliana* Columbia-0 (Col-0) seedlings to the respective half-maximal effect concentration (EC_50_) of either compound (3.5 µM for APO [25] and 4 µM for momilactone B; Supplemental Figure 1). We sampled plant material for RNA extraction and RNA-seq library preparation after 1 h, 6 h, and 24 h of treatment (Supplemental Table 1). As control, we treated seedlings with the equivalent concentration of solvent (dimethyl sulfoxide, DMSO). Sequencing reads were mapped to the *A. thaliana* TAIR10 reference genome (arabidopsis.org) [26]. In a principal component analysis (PCA) of read counts of the 1,000 genes with the highest variance after variance stabilizing transformation using the *DESeq2* package [27], samples clustered by time point; treated groups separated from untreated ones, indicating transcriptional responses to either allelochemical (Figure 1A). To investigate which genes were differentially expressed in response to the treatments, we performed differential expression analysis relative to the untreated 0 h baseline sample and combined this with clustering of genes using weighted gene correlation network analysis (WGCNA) [28] (Figure 1B). For APO-treated samples, we observed three clusters (A3, A6, and A7) that contained genes that were more strongly up-regulated in APO than in control treatments. For the momilactone B-treated samples, the genes contained in cluster M1 were up-regulated upon momilactone B treatment.

**Figure 1:**
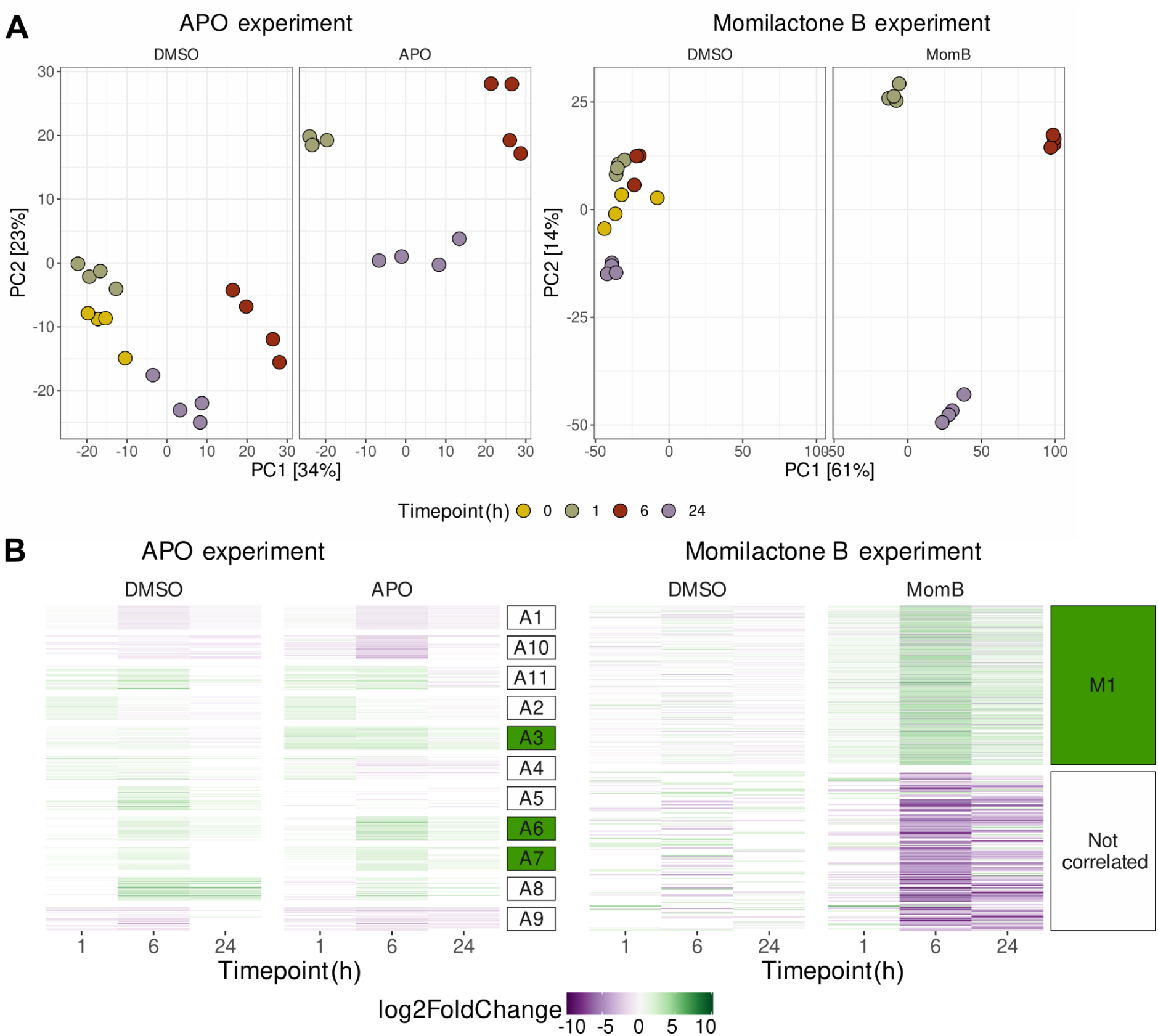
Allelochemicals elicit overlapping transcriptional changes in *A. thaliana*. **A)** Principal component analysis (PCA) of APO- and momilactone B-treated seedlings and controls. In the APO dataset, PC1 separates timepoints and PC2 separates treatments, while it is the other way around in the momilactone B dataset. **B)** Clustering of differentially expressed genes (DEGs). Heatmap of DEGs compared to the 0 h timepoint. Genes were clustered using WGCNA [28], the resulting clusters are indicated on the right. Clusters containing genes that were more strongly up-regulated in the treated than in the control samples are highlighted in green (A3 and A7 for APO and M1 for Momilactone B).

We proceeded to investigate if the up-regulated genes were involved in specific metabolic or cellular pathways and performed over-representation analysis (ORA) of the genes contained in the clusters A3, A6, A7, and M1, respectively, using the publicly available GOslim annotation of *A. thaliana* (arabidopsis.org) (Figure 2). We found that both treatments up-regulated genes involved in the activation of xenobiotics, such as e.g. cytochromes, oxidoreductases, or in glutathione-mediated detoxification, glycosylation, and transport, which are pillars of the three-phase xenobiotic detoxification system.

**Figure 2.**
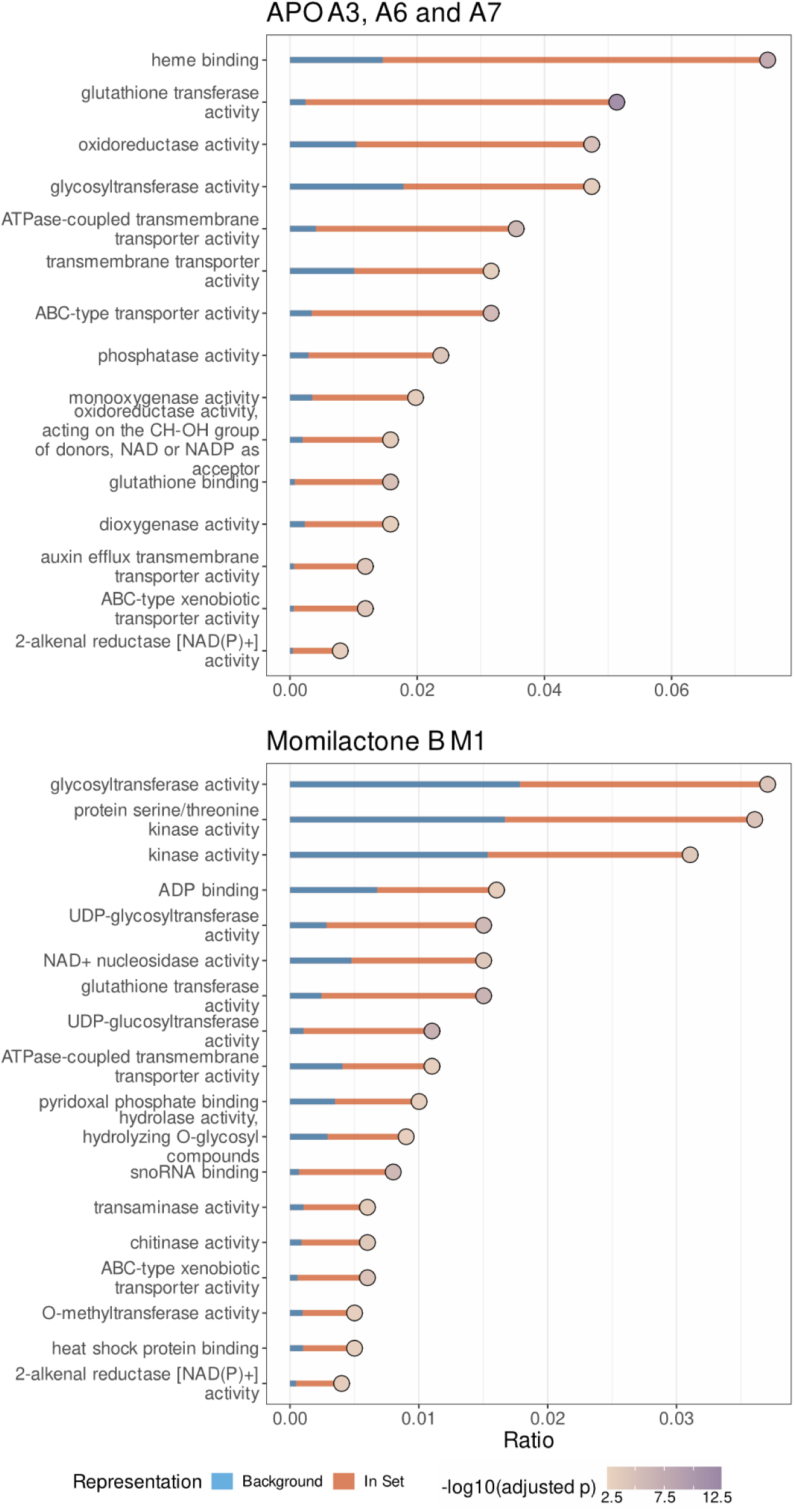
Gene ontology (GO)-term enrichment analysis of genes up-regulated by exposure to allelochemicals. GO-term analysis was performed on genes contained in clusters A3, A6 and A7 or M1 (see Figure 1B). Orange bars indicate the relative fraction of genes associated with the respective GO-term among all genes in the respective cluster; blue bars indicate the relative fraction of these genes among all genes in the genome. Redundant terms were removed. Fill color of the dots indicates -log_10_(p) of the hypergeometric test, adjusted using the method of Benjamini-Hochberg [29]. Only GO-terms that were significantly (p < 0.05) enriched are shown.

### General detoxification pathway genes are up-regulated upon allelochemical treatment

We next asked if the two allelochemicals simply activated the same general detoxification response, or if there was actual overlap among the differentially expressed genes. To this end, we performed an in-depth analysis of the genes involved in the different phases of the detoxification process: cytochrome P450 oxidases (CYPs) (phase I), glutathione-S-transferases (GSTs) and UPD-dependent glycosyltransferases (UGTs) (phase II), and transporters (phase III).

CYPs play a well-established role in the metabolic activation of xenobiotic compounds in phase I of the detoxification [30]. In our data, four *CYP* genes were up-regulated in both the APO and the momilactone B treatment, while eleven *CYP* genes were up-regulated only in APO and another eleven only in momilactone B treatment (Figure 3 and Supplemental Table 2). The up-regulated genes were distributed among all CYP clans. Noticeably, both treatments up-regulated genes from the *CYP81* family; *CYP81D8* and *CYP81D11* were up-regulated in both treatments, while other members were up-regulated upon either individual treatment (Figure 3 and Supplemental Table 2).

**Figure 3.**
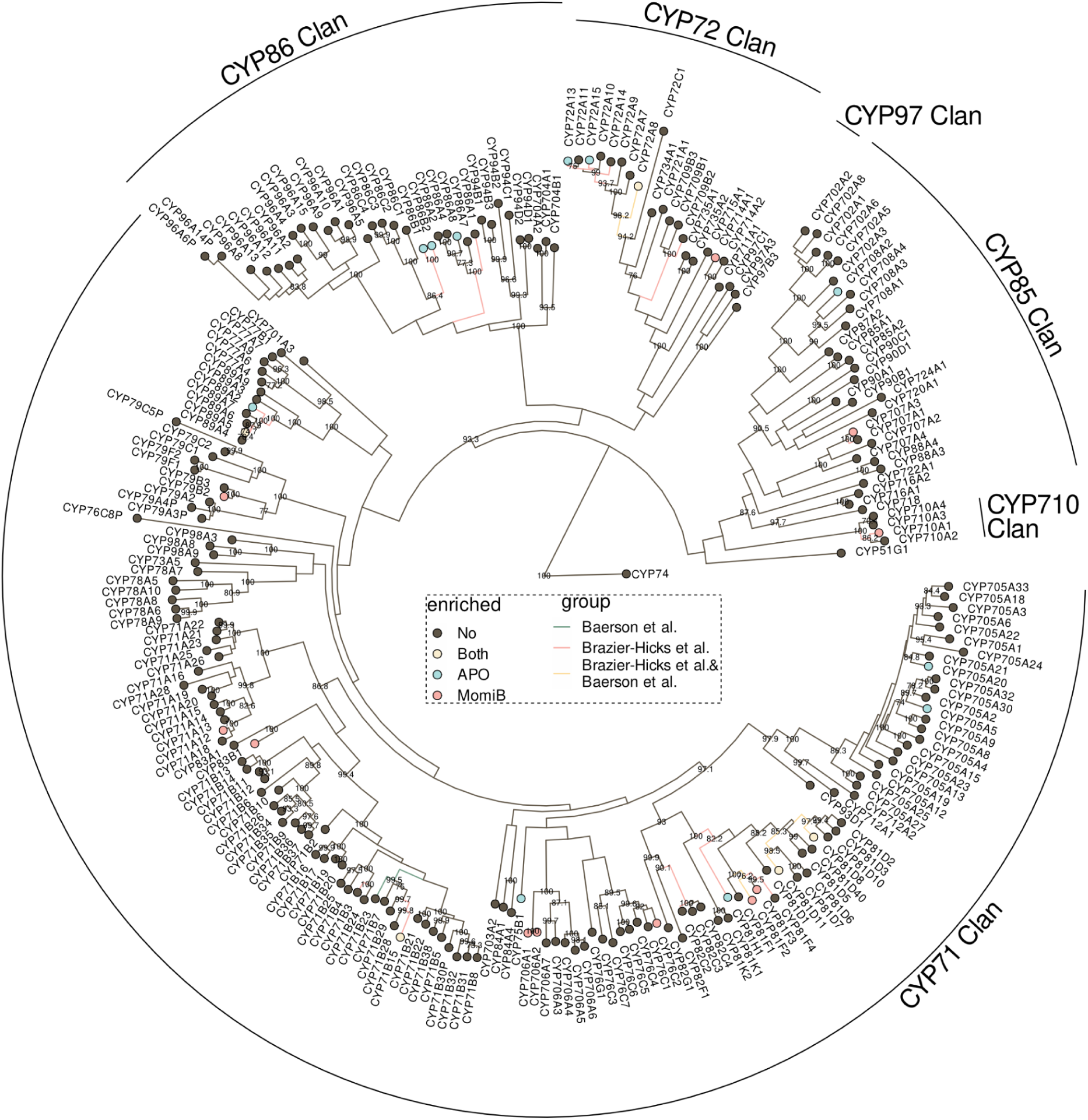
Up-regulation of *A. thaliana* cytochrome P450 oxidases (CYPs). Phylogenetic tree of all CYPs in the *A. thaliana* genome, based on protein sequence. Bootstrap values >70 are shown. CYP74 was used to root the tree. Coloured nodes indicate CYPs up-regulated by APO (blue), momilactone B (red) or both (yellow), coloured edges indicate CYPs up-regulated by BOA (blue; Baerson *et al*., 2005 [32]), fenclorim (red; Brazier-Hicks *et al*., 2018 [31]), or both (yellow).

To further understand the degree to which the the observed up-regulation was compound-specific, we added public information on the *A. thaliana* transcriptome response to the herbicide safener fenclorim [31] and to 2-benzoxazolinone (BOA), an intermediate in the bioconversion of DIBOA to APO [32]. Several of the genes up-regulated by APO and/or momilactone B were also up-regulated upon fenclorim or BOA treatment (Figure 3).Phase II detoxification involves the conjugation of sugar or glutathione to the activated compounds. Our initial analysis had already identified several *UGTs* and *GSTs* as being significantly up-regulated upon allelochemical treatment (Figure 1C). Multiple *UGTs* were up-regulated by both treatments (Figure 4A), with many belonging to the *UGT73* family. While most members of this family were up-regulated upon momilactone B treatment and by fenclorim [31], APO and its precursor BOA up-regulated only members of the *UGT73B* family. In addition to *UGT73s*, a few other *UGTs* from different families were up-regulated upon momilactone B treatment, many of these overlapping with *UGTs* up-regulated by herbicide safener treatment (Figure 4A) [31].

**Figure 4.**
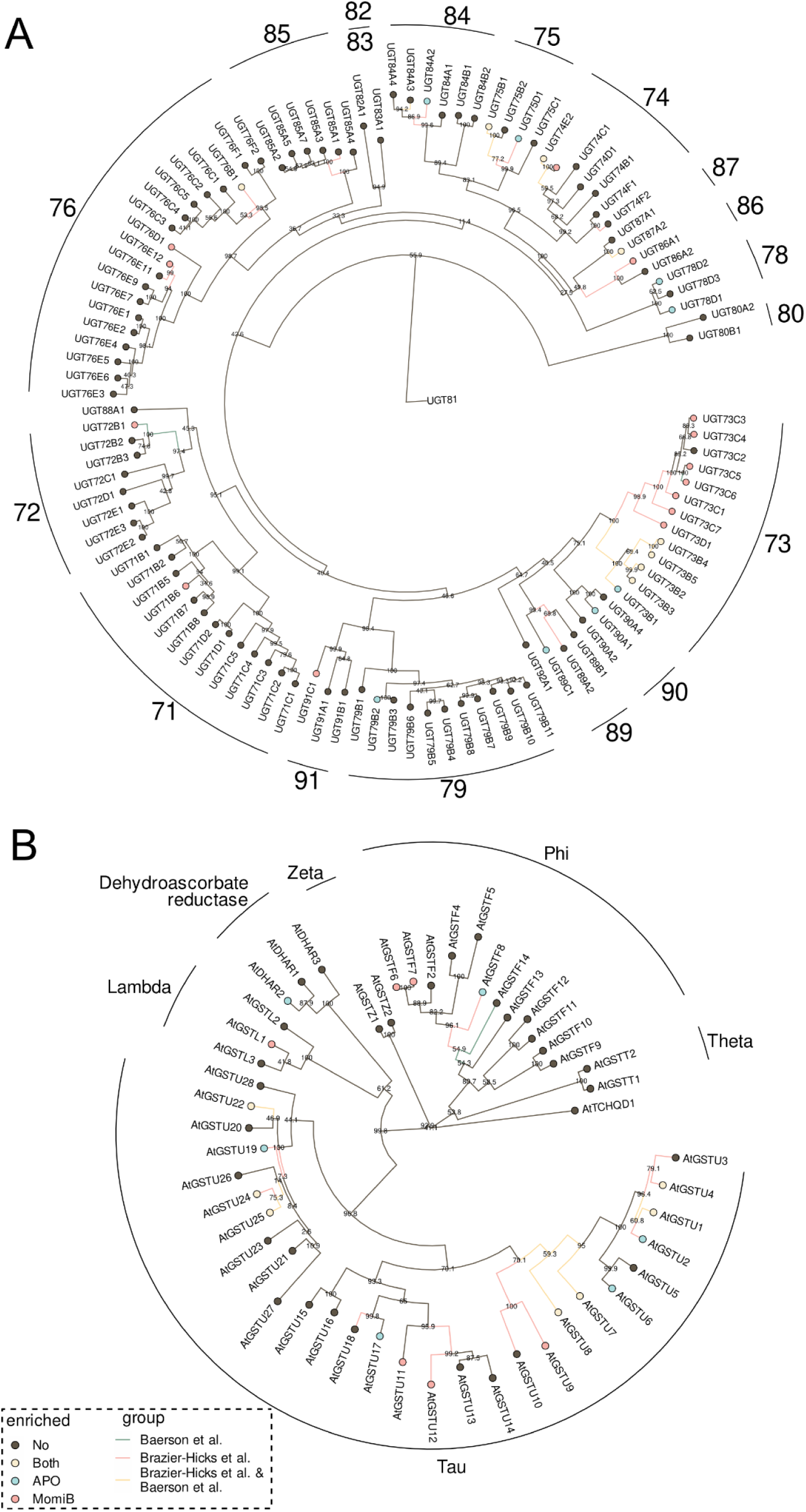
Up-regulation of *A*.*thaliana* UPD-dependent glycosyltransferases (UGTs) and glutathione-S-transferases (GSTs). Phylogenetic trees of all UGTs **(A)** and GSTs **(B)** in the *A. thaliana* genome, based on protein sequences. Coloured nodes indicate UGTs **(A)** and GSTs **(B)** up-regulated by APO (blue), momilactone B (red) or both (yellow), coloured edges indicate UGTs **(A)** and GSTs **(B)** up-regulated by BOA (blue; Baerson *et al*., 2005 [32]), fenclorim (red; Brazier-Hicks *et al*., 2018 [31]), or both (yellow).

Similar to the *UGTs*, several of the up-regulated *GSTs* overlapped between the APO and momilactone B treatments (Figure 4B). *GSTUs 1, 4, 7, 8, 22, 24*, and *25* were up-regulated by both APO and momilactone B, while tau class *GSTUs 9, 11*, and *12*, phi class *GSTs GSTF 6*, and *7*, and lambda class *GSTL1* were up-regulated only in response to momilactone B. Up-regulated in response to APO were tau class *GSTUs 2, 6, 17*, and *19, GSTF8* from the phi class, and *DHAR2* from the dehydroascorbate reductase clade. Most of the shared GSTs were also found to be up-regulated in response to fenclorim and BOA (Figure 4B).

The third and final phase of the detoxification process involves transport of the conjugated compounds into the vacuole or the apoplast. We found that treatment with APO and momilactone B both up-regulated transcripts coding for ABC-type transporters (Table 1), namely *AT2G47000, AT1G02520, AT3G47780*, and *AT3G59140*, all of which were reported to be up-regulated in the general detoxification process [33], which might suggest the sequestration of conjugated forms of these compounds into the vacuole or the apoplast.

**Table 1.**
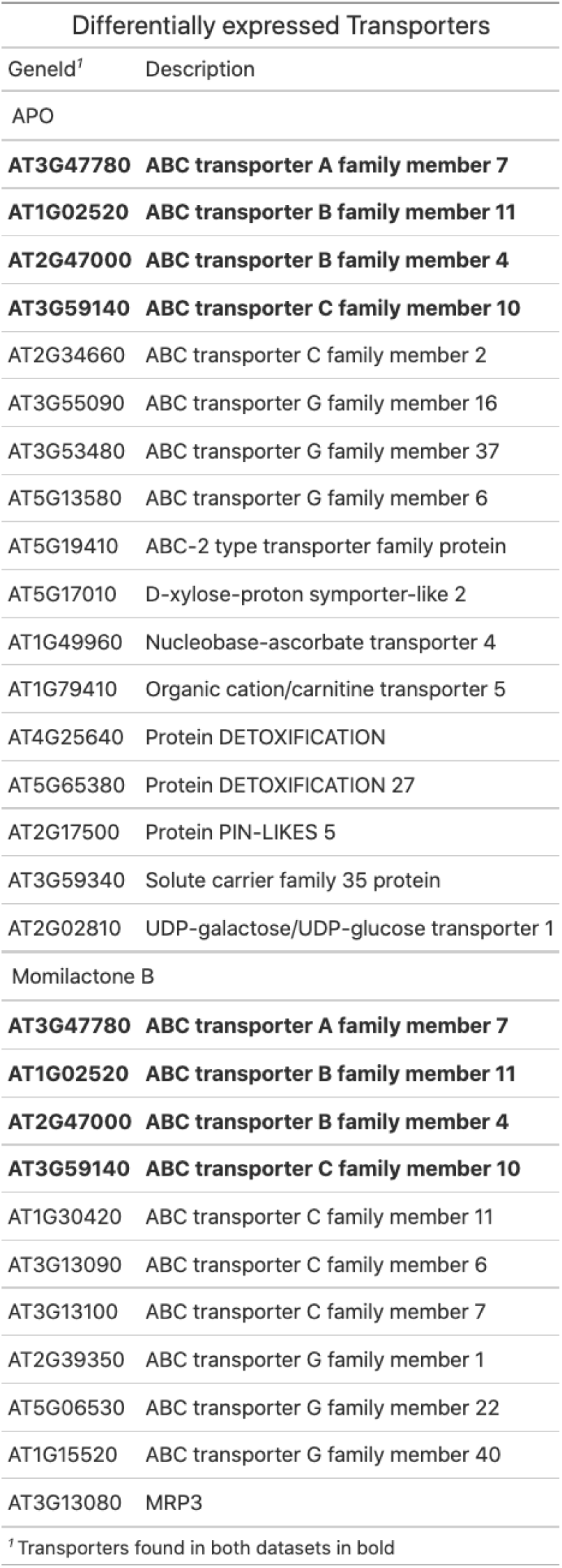
Transporters up-regulated by APO or momilactone B treatment. Marked in bold are transporters up-regulated in both treatments.

| Differentially expressed Transporters |  |
| --- | --- |
| GeneId<sup>†</sup> | Description |
| APO |  |
| **AT3G47780** | **ABC transporter A family member 7** |
| **AT1G02520** | **ABC transporter B family member 11** |
| **AT2G47000** | **ABC transporter B family member 4** |
| **AT3G59140** | **ABC transporter C family member 10** |
| AT2G34660 | ABC transporter C family member 2 |
| AT3G55090 | ABC transporter G family member 16 |
| AT3G53480 | ABC transporter G family member 37 |
| AT5G13580 | ABC transporter G family member 6 |
| AT5G19410 | ABC-2 type transporter family protein |
| AT5G17010 | D-xylose-proton symporter-like 2 |
| AT1G49960 | Nucleobase-ascorbate transporter 4 |
| AT1G79410 | Organic cation/carnitine transporter 5 |
| AT4G25640 | Protein DETOXIFICATION |
| AT5G65380 | Protein DETOXIFICATION 27 |
| AT2G17500 | Protein PIN-LIKES 5 |
| AT3G59340 | Solute carrier family 35 protein |
| AT2G02810 | UDP-galactose/UDP-glucose transporter 1 |
| Momilactone B |  |
| **AT3G47780** | **ABC transporter A family member 7** |
| **AT1G02520** | **ABC transporter B family member 11** |
| **AT2G47000** | **ABC transporter B family member 4** |
| **AT3G59140** | **ABC transporter C family member 10** |
| AT1G30420 | ABC transporter C family member 11 |
| AT3G13090 | ABC transporter C family member 6 |
| AT3G13100 | ABC transporter C family member 7 |
| AT2G39350 | ABC transporter G family member 1 |
| AT5G06530 | ABC transporter G family member 22 |
| AT1G15520 | ABC transporter G family member 40 |
| AT3G13080 | MRP3 |
| <sup>†</sup> Transporters found in both datasets in bold |  |

### Genes encoding plant cell wall constituents are downregulated in response to allelochemicals

Downregulated genes in cluster A10 were enriched for genes involved in cell wall integrity (structural constituent of cell wall and xyloglucan glycosylases) or in the peroxisome (heme binding and purine transport). The downregulated genes not assigned to a cluster in the momilactone B samples contained hydrolases and glycosylases (Supplemental Figure 2).

## Discussion

We treated *A. thaliana* with the two chemically very distinct, agriculturally relevant allelochemicals APO and momilactone B to understand the transcriptional response to these compounds. Both compounds are produced by cereal crops; BX are produced by wheat, maize and rye, while momilactones are produced by rice. The genes involved in both pathways are localized in biosynthetic gene clusters [17, 34]. Compounds appear to be produced in a species-specific manner; to date, no species has been identified that is capable of producing both classes of allelochemicals. The genome of *Echinochloa crus-galli*, a notorious weed in rice fields, contains biosynthetic gene clusters for both BX and momilactone A (the precursor of momilactone B) [35]. However, neither has the presence of momilactones in *E. crus-galli* been reported in the literature nor were we able to detect them by liquid chromatography–mass spectrometry (LC–MS) analysis (Supplemental Figure 3). That these two biosynthetic pathways seem to be mutually exclusive in grasses may indicate that grasses produce either BX or momilactones, but not both. One can therefore speculate that the two compound classes fulfill similar biological functions, and that there is no benefit in producing both. Although the two compounds have very different chemical structures, there was a significant overlap of genes up-regulated in response to both treatments, noticeably of genes that could be attributed the general detoxification response. The fact that up-regulated genes in our dataset overlapped with those from two published *A*.*thaliana* transcriptomic analyses in response to the herbicide safener fenclorim [31] and the BX BOA [32] further supported the notion that allelochemicals evoke the general detoxification response.

Among the genes up-regulated by either treatment or by both, UGTs, CYPs, and GSTs were overrepresented. One ongoing question about especially UGTs and CYPs involved in detoxification of herbicides and xenobiotics is whether their activity is specific for detoxification or if they usually have a different *in planta* role but are promiscuous and thus can act on both endogenous and exogenous substrates. With more genes from these families being characterized and their functions being identified, evidence points towards the latter [36, 37]. Our data further support this notion, even though further investigations are necessary to show that up-regulation of these genes in response to BX and momilactone B is accompanied by the according chemical modifications to these substrates.

CYPs involved in diterpene metabolism typically belong to the CYP71 and CYP85 clans; some can be found in the CYP72 clan [38], including those that were up-regulated by momilactone B.Momilactone B also up-regulated expression of *CYP710A1* from the CYP710 clan. Only few CYPs involved in diterpenoid metabolism have been identified in Arabidopsis, and the vast majority of CYP functions is still unknown. *CYP708A2* involved in thalianol to 7ß-hydroxythalianol biosynthesis [39] was up-regulated upon APO treatment, while several CYPs involved in camalexin and glucosinolate biosynthesis [40, 41] were enriched in the momilactone B up-regulated modules. This supports potentially significant enzyme promiscuity among CYPs that could explain their role in xenobiotic detoxification. The CYPs induced by both APO and momilactone B, as well as by fenclorim and BOA were all CYP81 family members. CYP81s have been linked to stress resistance and xenobiotic detoxification in *A. thaliana* and other plants [31–33, 42, 43].

*UGT73Bs* are consistently up-regulated upon allelochemical treatment (Figure 4). *UGT73B3* and *UGT73B5* are also involved in *A. thaliana* redox response to pathogenic Pseudomonas pathogens [44, 45]. *UGT73Bs* up-regulated upon fenclorim treatment were shown to glycosylate a number of different xenobiotics *in vitro*, indicating that these *UGTs* play a direct role in protection against such compounds [31]. Momilactone B and fenclorim treatments also up-regulated *UGT73Cs*; among these, *UGT73C5* encodes a known brassinosteroid-O-glycosyltransferase [37] that can also glycosylate and detoxify the mycotoxin deoxynivalenol [46], while *UGT73C6* encodes a UDP-glucose:flavonol-3-O-glycoside-7-O-glucosyltransferase [47].

Momilactone B is a diterpenoid, so we expected UGTs known to glycosylate diterpenoids to be up-regulated. These include UGTs from families 73, 74, 75, 76 and 85 [48]. Apart from the UGT73s mentioned above, several of the genes up-regulated by momilactone B belong to this UGT family. However, many of these UGTs were also induced by fenclorim treatment [31] (figure UGTs), which means that their transcriptional activation is most likely not a diterpenoid-specific response.

The majority of GSTs up-regulated in our study belong to the plant specific tau- and phi-classes that are known to be responsible for glutathione mediated detoxification of xenobiotics [49]. Of the seven tau-class GSTs (*GSTU1, 4, 7, 8, 22, 24* and *25*) up-regulated by both APO and momilactone B treatment, all except *GSTU4, 9* and *24* were also activated by fenclorim and BOA treatment (Figure 4B), supporting a robust detoxification response of these GSTs. Arabidopsis tau- and phi-class GSTs heterologously expressed in *E. coli* [50, 51] and in yeast [52] showed widespread ability to conjugate different herbicides to GSH. GSTLs and DHAR contain a cysteine instead of the active serine residue usually found in the catalytic site of GSTs, and presumably have no GSH conjugating activity [53], but can be involved in recycling specialized metabolites [54]. *GSTL1* and *DHAR2* were up-regulated upon momilactone B, and upon both momilactone B and APO treatment, respectively. Since they likely do not conjugate GSH, they may rather be involved in recycling GSH [55] in response to allelochemical treatment. GSTs have been known to be up-regulated by xenobiotics without playing a direct role in their conjugation [56], and part of the GST response to allelochemicals may be based on their general antioxidant function related to environmental stress [57]. Whether the GSTs up-regulated in our data conjugate allelochemicals to GSH therefore remains to be determined.

The final step in the three-phase detoxification is the transport of conjugated compounds into the vacuole or apoplast. All of the four ABC transporters up-regulated by both APO and momilactone B treatment were reported to be up-regulated in the general detoxification process [33], and *AT2g4700* and *AT3G59140* were further found to be up-regulated in response to methanol toxicity [58] and BOA treatment [32]. Since their discovery in plants [59], the MRP-class of ABC transporters has been associated with transport of GSH-conjugated xenobiotics [60], although MRP transporters are now known to transport a variety of substrates [61]. In our data, *MRP14* (*AT3G59140*) was up-regulated in response to both APO and momilactone B treatment; *MRP2* (*AT2g34660*) was up-regulated by APO, and *MRP12 (AT1G30420), MRP9 (AT3G13090), MRP7 (AT3G13100)*, and *MRP3 (AT3g13080)* by momilactone B. MRP3 transports GSH-conjugated xenobiotics and chlorophyll catabolites [62], but no specific transport activity has been reported for the other five MRP transporters. Nonetheless, the transcriptional activation of MRP transporters supports our notion that allelochemicals, like herbicides, are detoxified through the three-phase detoxification system.

## Conclusion

In summary, our data show that chemically diverse, phytotoxic compounds that are employed in inhibitory plant-plant interaction can trigger similar detoxification responses in the target plant, and provide insights into possible mechanisms of allelochemical tolerance. Further studies are necessary to investigate the metabolic processes and the chemical nature of potential conjugates. The detoxification of possibly conjugated allelochemicals by plant transporters may also explain how herbicide-responsive detoxification systems are maintained in plant populations that are typically not exposed to synthetic herbicides: since plants have to cope with a wide array of compounds released by other organisms, they have evolved and retained an arsenal of promiscuous enzymes that are able to detoxify harmful molecules with a certain degree of agnosticism towards their origin.

## Material and Methods

### Sample preparations and transcriptome sequencing

*A. thaliana* grown hydroponically in ½ MS media (pH 5.8) without sugar for 3 weeks (APO) or 6 days (momilactone B) were treated with either 3.5 µM APO, 4 µM momilactone B, or the equivalent concentration of the solvent DMSO. Tissue from four biological replicates per treatment (minimum 20 pooled seedlings per replicate) was collected 1, 6, and 24 h after beginning of the treatment and flash-frozen in liquid nitrogen. The tissue was lysed using a Retsch Mill (Retsch) before extracting total RNA using the RNeasy Plant Mini kit (Qiagen). RNA yield was measured using Qubit (Invitrogen) and RNA integrity was confirmed by agarose gel electrophoresis. Barcoded mRNA libraries were generated using either TruSeq RNA Sample Prep kit v2 (Illumina) (APO) or NEBNext RNA Ultra II Directional Library Kit (New England Biolabs) (momilactone B) following the manufacturers’ instructions. Libraries were sequenced on an Illumina HiSeqV4 as 100 bp single-end reads (APO) or an Illumina NovaSeq as 150 bp paired-end reads (momilactone B).

### Read mapping and quantification

All reads were mapped to the *A. thaliana* Col-0 TAIR10 reference genome (arabidopsis.org) (Supplemental Table 1). Mapping and feature counting were done using the nf-core pipeline RNAseq v3.5 [63, 64] with default parameters. In brief, reads were mapped using STAR and quantified using salmon [65, 66]. The counts per gene were further analyzed using R v4.1.2.

### Data analysis

Data analysis was carried out using R v4.1.2 and is documented in (https://github.com/nschan/Knoch_et_al_transcriptomes). In brief, count tables were imported into R. Differential expression analysis was carried out using *DESeq2* [27], weighted gene correlation network analysis was performed using *WGCNA* [28], and the beta parameter for WGCNA was picked automatically using *CEMiTool* [67]. Over-representation analysis of GO terms was performed using the enricher function from the *clusterProfiler* package [68]. Only genes that were contained in the clusters A3, A6, A7 or M1 and had a log2FoldChange > 0 and adjusted p-value below 0.01 at any time-point in the differential analysis were included in the overrepresentation analysis for upregulated genes, while the analysis of downregulated genes included genes contained in cluster A10, or not assigned to a cluster but included in the WGCNA analysis (momilactone B) with a log2FoldChange < 0 and an adjusted p-value below 0.01 in the differential expression analysis.

For phylogenetic analysis, protein sequences were aligned using the *AlignSeqs* function from *DECIPHER* [69], adjusted using the *AdjustAlignment* function, and maximum likelihood trees were fitted and bootstrapped 1000 times using the *phangorn* package [70] using the WAG substitution matrix.

### Momilactone D dose response

*A. thaliana* Col-0 seeds were sterilized with chlorine gas for 1 h and stratified for 6 d in the dark at 4°C, then sown on half strength Murashige and Skoog (MS) media supplemented with various concentrations of momilactone B dissolved in DMSO, to a final DMSO concentration of 0.1% in all plates. Seedlings were grown in a 16 h/8 h light/dark cycle chamber at 21°C with a light intensity of 50 µM/m2/sec. After 5 d of growth, the seedlings were imaged using a fixed camera and a ruler for scale. The primary root length was traced using ImageJ and primary root length was calculated using the scale as a reference. Primary root length was plotted as relative percentage of growth compared to the control sample. The drc package in RStudio [71] was used to fit a dose-response model and to calculate the half-maximal-effect concentration; data was plotted using the ggplot2 package [72].

### Momilactone content in Oryza sativa cv. Kitaake and Echinochloa crus-galli

*Oryza sativa* cv. Kitaake and *Echinochloa crus-galli* were grown in sand in the greenhouse for three weeks. Roots were harvested, weighed, snap-frozen in liquid nitrogen and homogenized. Metabolites were extracted in 10 times MeOH to sample weight, and momilactone A and B were measured on a Vanquish HPLC (Thermo Fisher Scientific) coupled via electrospray ionization to an TSQ Altis (Thermo Fisher Scientific) mass spectrometer, and quantified using authentic standards provided by Kazunori Okada, University of Tokyo.

## Data availability

Sequencing reads have been deposited in the ENA Short Read Archive (www.ebi.ac.uk/ena/) under study accession number PRJEB51016.

## Acknowledgements

We would like to thank the Next-Generation Sequencing unit at the Vienna Biocenter Core Facilities (VBCF) for their help in generating the sequencing data, Kazunori Okada, University of Tokyo, for providing momilactone A and B, and Katharina Jandrasits for technical lab support. LC-MS/MS analysis was performed by the Metabolomics Facility at Vienna BioCenter Core Facilities (VBCF), a member of the Vienna BioCenter (VBC), Austria, and funded by the City of Vienna through the Vienna Business Agency.

## Funding

This research was supported by the Austrian Academy of Sciences (ÖAW) via funding to C.B. and via a DOC fellowship to N.S.S., grant ID 25652; by the Austrian Science Fund (FWF) via a Lise Meitner Fellowship grant to E.K., grant ID FWF 2484-B21; and by the European Union’s Horizon 2020 research and innovation programme by the European Research Council (ERC), Grant Agreement No. 716823 ‘FEAR-SAP’.

## Author contributions

CB and NS conceived the study; EK, JK, SD, RS, and NSS performed the experiments; EK, NS, JK, CB, RS, and SD analyzed the data; EK, CB, and NS wrote the manuscript.

## Declaration of conflict of interest

The authors declare no conflict of interest.

## Notes

https://github.com/nschan/Knoch_et_al_transcriptomes

### Competing Interest Statement

The authors have declared no competing interest.

