## Supplemental Figures and Tables for "Transcriptional response of a target plant to benzoxazinoid and diterpene allelochemicals highlights commonalities in detoxification"

This file contains:

- Supplemental Figures 1-3
- Supplemental Tables 1-2
- Supplemental References

Supplemental Figure 1

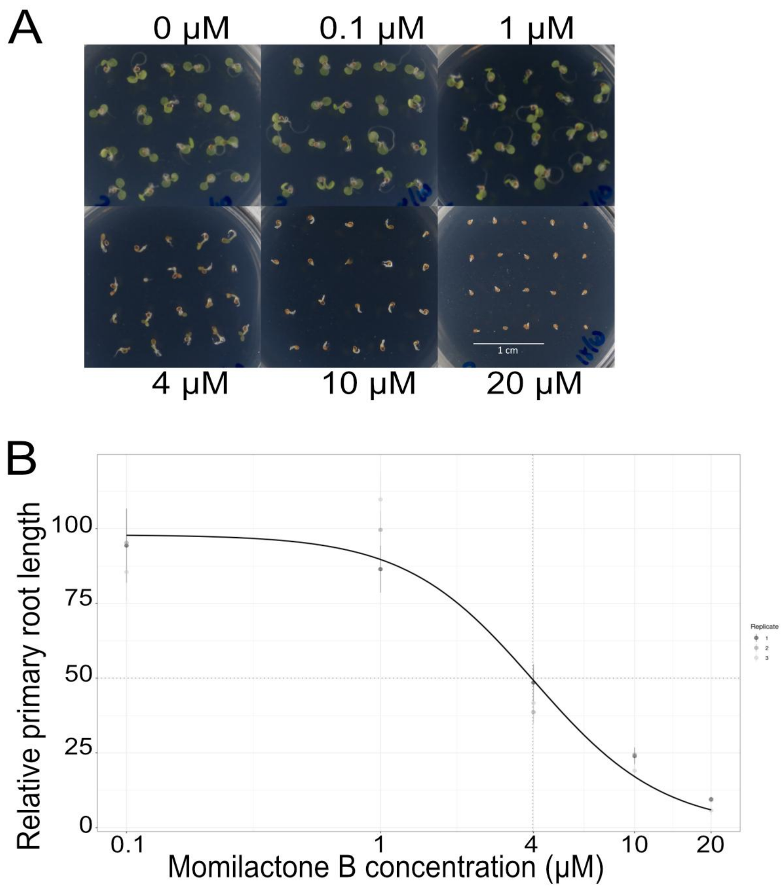


**Supplemental Figure 1. Momilactone B dose response**. **(A)** Growth phenotype of *A. thaliana* seedlings grown on ½ MS-agar with varying concentrations of momilactone B. Scale bar = 1 cm. **(B)** Primary root length plotted as relative percentage of growth compared to the control sample.

Supplemental Figure 2

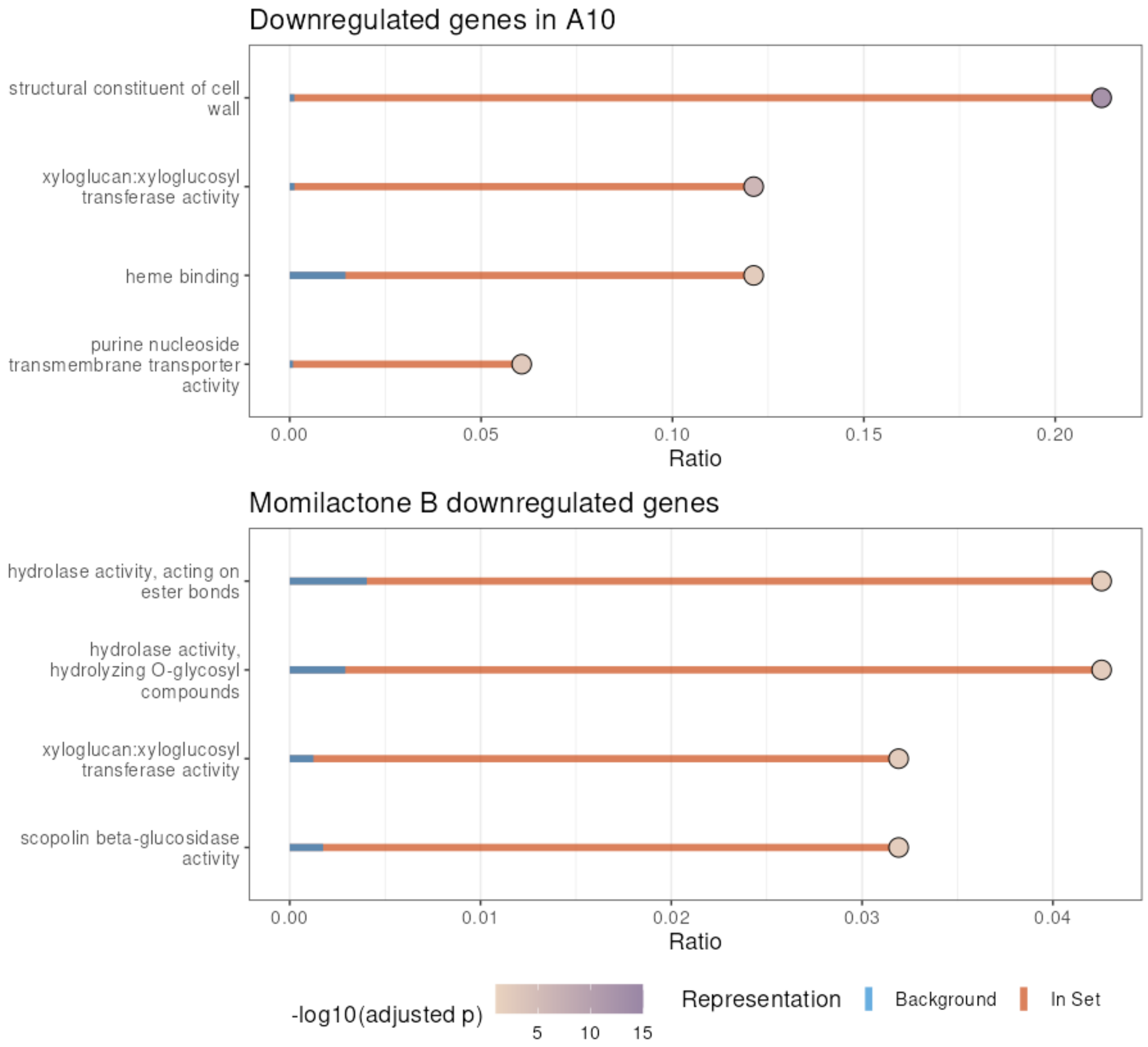


**Supplemental Figure 2: Overrepresentation analysis of down-regulated genes.** Lollipop plot of gene ontology (GO) terms of genes included in clusters A5 and A10 (APO) or not part of a cluster ("Not correlated", momilactone B). Genes with a negative log fold change and an adjusted p-value < 0.01 were included in an overrepresentation analysis of GO terms. Orange bars indicate the number of genes belonging to a particular GO-term relative to the total number of genes belonging to the term, blue bars indicate number of genes belonging to the GO-term compared to the total number of genes in the genome. Circle fill color indicates the p-value of the hypergeometric test, adjusted for multiple comparisons using the method of Benjamini-Hochberg [1].

Supplemental Figure 3

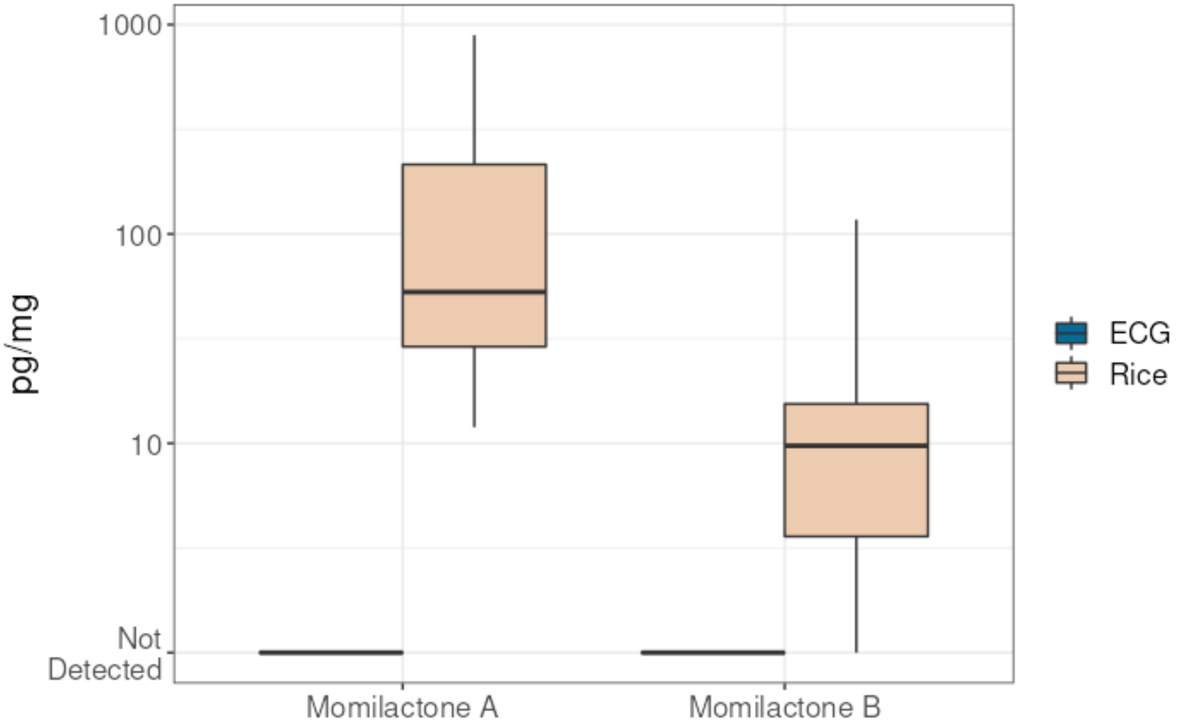


**Supplemental Figure 3. Momilactone content in Oryza sativa cv. Kitaake (Rice) and Echinochloa crus-galli (ECG).** Momilactone A and B in MeOH extracts from roots of three week old rice and *E. crus-galli* were measured by LCMS. Boxplots summarizing 18 replicates are shown, boxes indicate 1st to 3rd quartile, horizontal line indicates the median, whiskers extend to quartile1-1,5*IQR and quartile3+1,5*IQR

### Supplemental Table 1

| Sample | Experiment | Input | Unique mapped | % Unique mapped | Multimapped | % Multimapped |
|---|---|---|---|---|---|---|
| APO_1_A | APO | 33468367 | 32635808 | 97.51% | 598071 | 1.79% |
| APO_1_B | APO | 13303280 | 12958891 | 97.41% | 253520 | 1.91% |
| APO_1_C | APO | 13207791 | 12886546 | 97.57% | 226331 | 1.71% |
| APO_1_D | APO | 13831522 | 13492279 | 97.55% | 241370 | 1.75% |
| APO_24_A | APO | 12077780 | 11739344 | 97.20% | 237876 | 1.97% |
| APO_24_B | APO | 12844730 | 12472823 | 97.10% | 285025 | 2.22% |
| APO_24_C | APO | 13157352 | 12804481 | 97.32% | 267380 | 2.03% |
| APO_24_D | APO | 12244253 | 11917106 | 97.33% | 244987 | 2.00% |
| APO_6_A | APO | 15201412 | 14850778 | 97.69% | 247448 | 1.63% |
| APO_6_B | APO | 14864098 | 14484105 | 97.44% | 288045 | 1.94% |
| APO_6_C | APO | 12675530 | 12360557 | 97.52% | 234350 | 1.85% |
| APO_6_D | APO | 14154148 | 13800289 | 97.50% | 244854 | 1.73% |
| DMSO_0_A | APO | 18604718 | 18079070 | 97.17% | 379269 | 2.04% |
| DMSO_0_B | APO | 15526404 | 15082736 | 97.14% | 345099 | 2.22% |
| DMSO_0_C | APO | 14478446 | 14089113 | 97.31% | 282423 | 1.95% |
| DMSO_0_D | APO | 12652797 | 12331808 | 97.46% | 234548 | 1.85% |
| DMSO_1_A | APO | 21520940 | 20919612 | 97.21% | 450808 | 2.09% |
| DMSO_1_B | APO | 15108745 | 14695726 | 97.27% | 313665 | 2.08% |
| DMSO_1_C | APO | 12995297 | 12647493 | 97.32% | 250166 | 1.93% |
| DMSO_1_D | APO | 11787630 | 11471172 | 97.32% | 226384 | 1.92% |
| DMSO_24_A | APO | 14126302 | 13711086 | 97.06% | 322305 | 2.28% |
| DMSO_24_B | APO | 15213373 | 14818120 | 97.40% | 299031 | 1.97% |
| DMSO_24_C | APO | 12714549 | 12350707 | 97.14% | 278302 | 2.19% |
| DMSO_24_D | APO | 12464661 | 12107701 | 97.14% | 253034 | 2.03% |
| DMSO_6_A | APO | 14850544 | 13721149 | 92.39% | 817986 | 5.51% |
| DMSO_6_B | APO | 15272280 | 14626289 | 95.77% | 323714 | 2.12% |
| DMSO_6_C | APO | 13155897 | 12451840 | 94.65% | 212284 | 1.61% |
| DMSO_6_D | APO | 13227781 | 12609698 | 95.33% | 258050 | 1.95% |
| DMSO_0h_1 | Momilactone B | 14382197 | 1123045 | 7.81% | 81329 | 0.57% |
| DMSO_0h_2 | Momilactone B | 3958297 | 3469868 | 87.66% | 330821 | 8.36% |
| DMSO_0h_3 | Momilactone B | 1417280 | 780139 | 55.04% | 365062 | 25.76% |
| DMSO_0h_4 | Momilactone B | 10901026 | 9199207 | 84.39% | 1481899 | 13.59% |
| DMSO_1h_1 | Momilactone B | 10194188 | 8774305 | 86.07% | 978795 | 9.60% |
| DMSO_1h_2 | Momilactone B | 14674393 | 13351529 | 90.99% | 945121 | 6.44% |
| DMSO_1h_3 | Momilactone B | 18902989 | 14669222 | 77.60% | 2630303 | 13.91% |
| DMSO_1h_4 | Momilactone B | 9533524 | 8233576 | 86.36% | 843176 | 8.84% |
| DMSO_24h_1 | Momilactone B | 22178704 | 20128771 | 90.76% | 1267084 | 5.71% |
| DMSO_24h_2 | Momilactone B | 24088746 | 20114579 | 83.50% | 3123586 | 12.97% |
| DMSO_24h_3 | Momilactone B | 20218885 | 17022949 | 84.19% | 2460316 | 12.17% |
| DMSO_24h_4 | Momilactone B | 23610363 | 21046925 | 89.14% | 1541774 | 6.53% |
| DMSO_6h_2 | Momilactone B | 21041416 | 18199018 | 86.49% | 1376492 | 6.54% |
| DMSO_6h_3 | Momilactone B | 22702394 | 19229227 | 84.70% | 2805472 | 12.36% |
| DMSO_6h_4 | Momilactone B | 24868740 | 18401963 | 74.00% | 5853970 | 23.54% |
| MomB_1h_1 | Momilactone B | 19396480 | 17997922 | 92.79% | 1152979 | 5.94% |
| MomB_1h_2 | Momilactone B | 15280820 | 12593194 | 82.41% | 2351135 | 15.39% |
| MomB_1h_3 | Momilactone B | 24227029 | 21875372 | 90.29% | 1885258 | 7.78% |
| MomB_1h_4 | Momilactone B | 19499176 | 15968805 | 81.89% | 3216737 | 16.50% |
| MomB_24h_1 | Momilactone B | 29161388 | 24142676 | 82.79% | 4103996 | 14.07% |
| MomB_24h_2 | Momilactone B | 22966052 | 19388213 | 84.42% | 2947973 | 12.84% |
| MomB_24h_3 | Momilactone B | 15904026 | 14125386 | 88.82% | 1309675 | 8.23% |
| MomB_24h_4 | Momilactone B | 16506046 | 14677116 | 88.92% | 1517243 | 9.19% |
| MomB_6h_1 | Momilactone B | 16784399 | 15841123 | 94.38% | 781481 | 4.66% |
| MomB_6h_2 | Momilactone B | 17026600 | 15717196 | 92.31% | 1053980 | 6.19% |
| MomB_6h_3 | Momilactone B | 24497494 | 21802140 | 89.00% | 2115931 | 8.64% |
| MomB_6h_4 | Momilactone B | 13566887 | 11009230 | 81.15% | 2273091 | 16.75% |

**Supplemental Table 1. RNA-seq mapping statistics.**

### Supplemental Table 2

Supplemental Table 2: Differentially expressed CYP450s

| CYP | GeneId | Also found in |
|---|---|---|
| APO | | |
| CYP705A20 | AT3G20110 | - |
| CYP705A5 | AT5G47990 | - |
| CYP708A2 | AT5G48000 | - |
| CYP71B15 | AT3G26830 | Brazier-Hicks et al. |
| CYP72A13 | AT3G14660 | Brazier-Hicks et al. |
| CYP72A15 | AT3G14690 | Brazier-Hicks et al. |
| CYP72A8 | AT3G14620 | Brazier-Hicks et al. & Baerson et al. |
| CYP75B1 | AT5G07990 | - |
| CYP81D11 | AT3G28740 | Brazier-Hicks et al. & Baerson et al. |
| CYP81D8 | AT4G37370 | Brazier-Hicks et al. & Baerson et al. |
| CYP81H1 | AT4G37310 | Brazier-Hicks et al. |
| CYP86A8 | AT2G45970 | - |
| CYP86B1 | AT5G23190 | Brazier-Hicks et al. |
| CYP86B2 | AT5G08250 | Brazier-Hicks et al. |
| CYP89A2 | AT1G64900 | Brazier-Hicks et al. |
| CYP89A5 | AT1G64950 | - |
| Momilactone B | | |
| CYP706A1 | AT4G22690 | - |
| CYP706A2 | AT4G22710 | - |
| CYP707A3 | AT5G45340 | - |
| CYP710A1 | AT2G34500 | Brazier-Hicks et al. |
| CYP714A1 | AT5G24910 | - |
| CYP71A12 | AT2G30750 | - |
| CYP71B15 | AT3G26830 | Brazier-Hicks et al. |
| CYP72A8 | AT3G14620 | Brazier-Hicks et al. & Baerson et al. |
| CYP76C2 | AT2G45570 | - |
| CYP79B2 | AT4G39950 | - |
| CYP81D11 | AT3G28740 | Brazier-Hicks et al. & Baerson et al. |
| CYP81D8 | AT4G37370 | Brazier-Hicks et al. & Baerson et al. |
| CYP81F2 | AT5G57220 | Brazier-Hicks et al. & Baerson et al. |
| CYP81F3 | AT4G37400 | - |
| CYP83B1 | AT4G31500 | - |
| CYP89A5 | AT1G64950 | - |

**Supplemental Table 2. Differentially expressed Cytochrome P450 oxidases (CYPs).** All CYPs that were significantly ($p < 0.01$) differentially expressed at any timepoint are listed. The first column provides the CYP name, the second column shows the corresponding Col-0 locus identifier, the last column indicates if the particular genes was also found as differentially expressed upon femclorin [2] or BOA treatment [3]. Bold type highlights CYPs that were differentially expressed upon both APO and momilactone treatment.
